# Nanopore sequencing enables near-complete *de novo* assembly of *Saccharomyces cerevisiae* reference strain CEN.PK113-7D

**DOI:** 10.1101/175984

**Authors:** Alex N. Salazar, Arthur R. Gorter de Vries, Marcel van den Broek, Melanie Wijsman, Pilar de la Torre Cortés, Anja Brickwedde, Nick Brouwers, Jean-Marc G. Daran, Thomas Abeel

## Abstract

The haploid *Saccharomyces cerevisiae* strain CEN.PK113-7D is a popular model system for metabolic engineering and systems biology research. Current genome assemblies are based on short-read sequencing data scaffolded based on homology to strain S288C. However, these assemblies contain large sequence gaps, particularly in subtelomeric regions, and the assumption of perfect homology to S288C for scaffolding introduces bias.

In this study, we obtained a near-complete genome assembly of CEN.PK113-7D using only Oxford Nanopore Technology’s MinION sequencing platform. 15 of the 16 chromosomes, the mitochondrial genome, and the 2-micron plasmid are assembled in single contigs and all but one chromosome starts or ends in a telomere cap. This improved genome assembly contains 770 Kbp of added sequence containing 248 gene annotations in comparison to the previous assembly of CEN.PK113-7D. Many of these genes encode functions determining fitness in specific growth conditions and are therefore highly relevant for various industrial applications. Furthermore, we discovered a translocation between chromosomes III and VIII which caused misidentification of a *MAL* locus in the previous CEN.PK113-7D assembly. This study demonstrates the power of long-read sequencing by providing a high-quality reference assembly and annotation of CEN.PK113-7D and places a caveat on assumed genome stability of microorganisms.

## Introduction

Whole Genome Sequencing (WGS) reveals important genetic information of an organism which can be linked to specific phenotypes and enable genetic engineering approaches (Mardis 2008, Ng and Kirkness 2010). Short-read sequencing has become the standard method for WGS in the past years due to its low cost, high sequencing accuracy and high output of sequence reads. In most cases, the obtained read data is used to reassemble the sequenced genome either by *de novo* assembly or by mapping the reads to a previously-assembled closely-related genome. However, the sequence reads obtained are relatively short: between 35 and 1000 bp (van Dijk *et al.* 2014). This poses challenges as genomes have long stretches of repetitive sequences of several thousand nucleotides in length and can only be characterized if a read spans the repetitive region and has a unique fit to the flanking ends (Matheson *et al.* 2017). As a result, *de novo* genome assembly based on short-read technologies “break” at repetitive regions preventing reconstruction of whole chromosomes. The resulting assembly consists of dozens to hundreds of sequence fragments, commonly referred to as *contigs*. These contigs are then either analysed independently or ordered and joined together adjacently based on their alignment to a closely-related reference genome. However, referenced based joining of contigs into so-called *scaffolds*, is based on the assumption that the genetic structure of the sequenced strain is identical to that of the reference genome—potentially concealing existing genetic variation.

Previous genome assemblies of the *Saccharomyces cerevisiae* strain CEN.PK113-7D have been based on homology with the fully-assembled reference genome of *S. cerevisiae* strain S288C (Cherry *et al.* 2012, Nijkamp *et al.* 2012). CEN.PK113-7D is a haploid strain used as a model organism in biotechnology-related research and systems biology because of its convenient growth characteristics, its robustness under industrially-relevant conditions, and its excellent genetic accessibility (Canelas *et al.* 2010, González-Ramos *et al.* 2016, Nijkamp *et al.* 2012, Papapetridis *et al.* 2017). CEN.PK113-7D was sequenced using a combination of 454 and Illumina short-read libraries and a draft genome was assembled consisting of over 700 contigs (Nijkamp *et al.* 2012). After scaffolding using MAIA (Nijkamp *et al.* 2010) and linking based on homology with the genome of S288C, it was possible to reconstruct all 16 chromosomes. However, there were large sequence gaps within chromosomes and the subtelomeric regions were left unassembled, both of which could contain relevant open reading frames (ORFs) (Nijkamp *et al.* 2012).Assuming homology to S288C, more than 90% of missing sequence was located in repetitive regions corresponding mostly to subtelomeric regions and Ty-elements. These regions are genetically unstable as repeated sequences promote recombination events (Pryde *et al.* 1995); therefore the assumption of homology with S288C could be unjustified. Ty-elements are present across the genome: repetitive sequences with varying length (on average ∼6 Kbp) resulting from introgressions of viral DNA (Kim *et al.* 1998). Subtelomeric regions are segments towards the end of chromosomes consisting of highly repetitive elements making them notoriously challenging to reconstruct using only short-read sequencing data (Bergström *et al.* 2014). While Ty-elements are likely to have limited impact on gene expression, subtelomeric regions harbour various so-called subtelomeric genes. Several gene families are present mostly in subtelomeric regions and typically have functions determining the cell’s interaction with its environment; such as nutrient uptake (Carlson *et al.* 1985, Naumov *et al.* 1995), sugar utilisation (Teste *et al.* 2010), and inhibitor tolerance (Denayrolles *et al.* 1997). Many of these subtelomeric gene families therefore contribute to the adaptation of industrial strains to the specific environment they are used in. For example, the *RTM* and *SUC* gene families are relevant for bioethanol production as they increase inhibitor-tolerance in molasses and utilization of extracellular sucrose, respectively (Carlson *et al.* 1985, Denayrolles *et al.* 1997). Similarly, *MAL* genes enable utilization of maltose and maltotriose and *FLO* genes enable calcium-dependent flocculation, both of which are crucial for the beer brewing industry (Brown *et al.* 2010, Lodolo *et al.* 2008, Teunissen and Steensma 1995). As is the case for Ty-elements, subtelomeric regions are unstable due to repetitive sequences and homology to various regions of the genome, which is likely to cause diversity across strains (Brown *et al.* 2010, Nijkamp *et al.* 2012, Pryde *et al.* 1995). Characterizing and accurately localizing subtelomeric gene families is thus crucial for associating strain performance to specific genomic features and for targeted engineering approaches for strain improvement (Bergström *et al.* 2014).

In contrast to short-read technologies, single-molecule sequencing technologies can output sequence reads of several thousand nucleotides in length. Recent developments of long-read sequencing technologies have decreased the cost and increased the accuracy and output, yielding near-complete assemblies of diverse yeast strains (Giordano *et al.* 2017, McIlwain *et al.* 2016). For example, *de novo* assembly of a biofuel production *S. cerevisiae* strain using PacBio reads produced a genome assembly consisting of 25 chromosomal contigs scaffolded into 16 chromosomes. This assembly revealed 92 new genes relative to S288C amongst which 28 previously uncharacterized and unnamed genes. Interestingly, many of these genes had functions linked to stress tolerance and carbon metabolism which are functions critical to the strains industrial application (McIlwain *et al.* 2016). In addition, rapid technological advances in nanopore sequencing have matured as a competitive long-read sequencing technology and the first yeast genomes assembled using nanopore reads are appearing (Giordano *et al.* 2017, Goodwin *et al.* 2015, Istace *et al.* 2017, Jansen *et al.* 2017, McIlwain *et al.* 2016). For example, Istace *et al.* sequenced 21 wild *S. cerevisiae* isolates and their genome assemblies ranged between 18 and 105 contigs enabling the detection of 29 translocations and 4 inversions relative to the chromosome structure of reference S288C. In addition, large variations were found in several difficult to sequence subtelomeric genes such as *CUP1*, which was correlated to large differences in copper tolerance (Istace *et al.* 2017). Nanopore sequencing has thus proven to be a potent technology for characterizing yeast.

In this study, we sequenced CEN.PK113-7D using Oxford Nanopore Technology’s (ONT) MinION sequencing platform. This nanopore *de nov*o assembly was compared to the previous short-read assembly of CEN.PK113-7D (Nijkamp *et al.* 2012) with particular attention for previously, poorly-assembled subtelomeric regions and for structural variation potentially concealed due to the assumption of homology to S288C.

## Materials and methods

### Yeast strains

The *Saccharomyces cerevisiae* strain “CEN.PK113-7D Frankfurt” (*MATa MAL2-8c*) was kindly provided by Dr. P. Kötter in 2016 (Entian and Kötter 2007, Nijkamp *et al.* 2012). It was plated on solid YPD (containing 10 g/l yeast extract, 20 g/l peptone and 20 g/l glucose) upon arrival and a single colony was grown once until stationary phase in liquid YPD medium and 1 mL aliquots with 30% glycerol were stored at −80°C since. The previously sequenced CEN.PK113-7D sample was renamed “CEN.PK113-7D Delft” (Nijkamp *et al.* 2012). It was obtained from the same source in 2001 and 1 mL aliquots with 30% glycerol were stored at −80°C with minimal propagation since (no more than three cultures on YPD as described above).

### Yeast cultivation and genomic DNA extraction

Yeast cultures were incubated in 500-ml shake-flasks containing 100 ml liquid YPD medium at 30°C on an orbital shaker set at 200 rpm until the strains reached stationary phase with an OD_660_ between 12 and 20. Genomic DNA of CEN.PK113-7D Delft and CEN.PK113-7D Frankfurt for whole genome sequencing was isolated using the Qiagen 100/G kit (Qiagen, Hilden, Germany) according to the manufacturer’s instructions and quantified using a Qubit^®^ Fluorometer 2.0 (ThermoFisher Scientific, Waltham, MA).

### Short-read Illumina sequencing

Genomic DNA of CEN.PK113-7D Frankfurt was sequenced on a HiSeq2500 sequencer (Illumina, San Diego, CA) with 150 bp paired-end reads using PCR-free library preparation by Novogene Bioinformatics Technology Co., Ltd (Yuen Long, Hong Kong). All Illumina sequencing data are available at NCBI (https://dwww.ncbi.nlm.nih.gov/) under the bioproject accession number PRJNA393501.

### MinION Sequencing

MinION genomic libraries were prepared using either nanopore Sequencing Kit SQK-MAP006 (2D-ligation for R7.3 chemistry), SQK-RAD001 (Rapid library prep kit for R9 chemistries) or SQK-MAP007 (2D-ligation for R9 chemistries) (Oxford Nanopore Technologies, Oxford, United Kingdom). Two separate libraries of SQK-MAP006 and one library of SQK-RAD001 were used to sequence CEN.PK113-7D Delft. Only one SQK-MAP007 library was used to sequence CEN.PK113-7D Frankfurt. With the exception of the SQK-RAD001 library, all libraries used 2-3 μg of genomic DNA fragmented in a Covaris g-tube (Covaris) with the “8-10 kbp fragments” settings according to manufacturer’s instructions. The SQK-RAD001 library used 200 ng of unsheared genomic DNA. Libraries for SQK-MAP006 and SQK-MAP007 were constructed following manufacturer’s instructions with the exception of using 0.4x concentration of AMPure XP Beads (Beckman Coulter Inc., Brea, CA) and 80% EtOH during the “End Repair/dA-tailing module” step. The SQK-RAD001 library was constructed following manufacturer’s instructions. Prior to sequencing, flow cell quality was assessed by running the MinKNOW platform QC (Oxford Nanopore Technology). All flow cells were primed with priming buffer and the libraries were loaded following manufacturer’s instructions. The mixture was then loaded into the flow cells for sequencing. The SQK-MAP006 library of CEN.PK113-7D Delft was sequenced twice on a R7.3 chemistry flow cell (FLO-MIN103) and the SQK-RAD001 library was sequenced on a R9 chemistry flow cell (FLO-MIN105)—all for 48 hours. The SQK-MAP007 library for CEN.PK113-7D Frankfurt was sequenced for 48 hours on a R9 chemistry flow cell (FLO-MIN104). Reads from all sequencing runs were uploaded and base-called using Metrichor desktop agent (https://metrichor.com/s/). The error rate of nanopore reads in the CEN.PK113-7D Frankfurt and Delft was determined by aligning them to the final CEN.PK113-7D assembly (see section below) using Graphmap (Sović *et al.* 2016) and calculating mismatches based on the CIGAR strings of reads with a mapping quality of at least 1 and no more than 500 nt of soft/hard clipping on each end of the alignment to avoid erroneous read-alignments due to repetitive regions (i.e. paralogous genes, genes with copy number variation). All nanopore sequencing data are available at NCBI under the bioproject accession number PRJNA393501.

### *De novo* genome assembly

FASTA and FASTQ files were extracted from base-called FAST5 files using Poretools (version 0.6.0) (Loman and Quinlan 2014). Raw nanopore reads were filtered for lambda DNA by aligning to the *Enterobacteria phage lambda* reference genome (RefSeq assembly accession: GCF_000840245.1) using Graphmap (Sović *et al.* 2016) with *–no-end2end* parameter and retaining only unmappeds reads using Samtools (Li *et al.* 2009). All reads obtained from the Delft and the Frankfurt CEN.PK113-7D stock cultures were assembled *de novo* using Canu (version 1.3) (Koren *et al.* 2017) with –*genomesize* set to 12 Mbp. The assemblies were aligned using the MUMmer tool package: Nucmer with the *–maxmatch* parameter and filtered for the best one-to-one alignment using Delta-filter (Kurtz *et al.* 2004). The genome assemblies were visualized using Mummerplot (Kurtz *et al.* 2004) with the *–fat* parameter. Gene annotations were performed using MAKER2 annotation pipeline (version 2.31.9) using SNAP (version 2013-11-29) and Augustus (version 3.2.3) as *ab initio* gene predictors (Holt and Yandell 2011). S288C EST and protein sequences were obtained from SGD (*Saccharomyces* Genome Database, http://www.yeastgenome.org/) and were aligned using BLASTX (BLAST version 2.2.28+) (Camacho *et al.* 2009). Translated protein sequence of the final gene model were aligned using BLASTP to S288C protein Swiss-Prot database. Custom made Perl scripts were used to map systematic names to the annotated gene names. Telomere cap sequences (TEL07R of size 7,306 bp and TEL07L of size 781 bp) from the manually-curated and complete reference genome for *S. cerevisiae* S288C (version R64, Genbank ID: 285798) obtained from SGD were aligned to the assembly as a proxy to assess completeness of each assembled chromosome. SGIDs for TEL07R and TEL07L are S000028960 and S000028887, respectively. The Tablet genome browser (Milne *et al.* 2012) was used to visualize nanopore reads aligned to the nanopore *de novo* assemblies. Short assembly errors in the Frankfurt assembly were corrected with Nanopolish (version 0.5.0) using default parameters (Loman *et al.* 2015). Two contigs, corresponding to chromosome XII, were manually scaffolded based on homology to S288C. To obtain the 2-micron native plasmid in CEN.PK113-7D, we aligned S288C’s native plasmid to the “unassembled” contigs file provided by Canu (Koren *et al.* 2017) and obtained the best aligned contig in terms of size and sequence similarity. Duplicated regions due to assembly difficulties in closing circular genomes were identified with Nucmer and manually corrected. BWA (Li and Durbin 2010) was used to align Illumina reads to the scaffolded Frankfurt assembly using default parameters. Pilon (Walker *et al.* 2014) was then used to further correct assembly errors by aligning Illumina reads to the scaffolded Frankfurt assembly using correction of only SNPs and short indels (*–fix bases* parameter) using only reads with a minimum mapping quality of 20 (–*minmq 20* parameter). Polishing with structural variant correction in addition to SNP and short indel correction was benchmarked, but not applied to the final assembly (Additional File 1).

### Analysis of added information in the CEN.PK113-7D nanopore assembly

Gained and lost sequence information in the nanopore assembly of CEN.PK113-7D was determined by comparing it to the previous short-read assembly (Nijkamp *et al.* 2012). Contigs of at least 1 Kbp of short-read assembly were aligned to the nanopore CEN.PK113-7D Frankfurt assembly using the MUMmer tool package (Kurtz *et al.* 2004) using *show-coords* to extract alignment coordinates. For multi-mapped contigs, overlapping alignments of the same contig were collapsed and the largest alignment length as determined by Nucmer was used. Unaligned coordinates in the nanopore assembly were extracted and considered as added sequence. Added genes were retrieved by extracting the gene annotations in these unaligned regions from the annotated nanopore genome; mitochondria and 2-micron plasmid genes were excluded For the lost sequence, unaligned sequences were obtained by aligning the contigs of the nanopore assembly to the short-read contigs of at least 1 kb using the same procedure as described above. Lost genes were retrieved by aligning the unaligned sequences to the short-read CEN.PK113-7D assembly with BLASTN (version 2.2.31+) (Camacho *et al.* 2009) and retrieving gene annotations. BLASTN was used to align DNA sequences of YHRCTy1-1, YDRCTy2-1, YILWTy3-1, YHLWTy4-1, and YCLWTy5-1 (obtained from the *Saccharomyces Genome Database;* SGIDs: S000007006, S000006862, S000007020, S000006991, and S000006831, respectively) as proxies for the location of two known groups of Ty-elements in *Saccharomyces cerevisiae*, *Metaviridae* and *Pseudoviridae (Kim et al. 1998)*, in the CEN.PK113-7D Frankfurt assembly. Non-redundant locations with at least a 2 Kbp alignment and an E-value of 0.0 as determined by BLASTN were then manually inspected.

### Comparison of the CEN.PK113-7D assembly to the S288C genome

The nanopore assembly of CEN.PK113-7D and the reference genome of S2888C (Accession number GCA_000146045.2) were annotated using the MAKER2 pipeline described in the “De novo genome assembly” section. For each genome a list of gene names per chromosome was constructed and compared strictly on their names to identify genes names absent in the corresponding chromosome in the other genome. The ORFs of genes identified as absent in either genome were aligned using BLASTN (version 2.2.31+) to the total set of ORFs of the other genome and matches with an alignment length of half the query and with a sequence identity of at least 95% were listed. If one of the unique genes aligned to an ORF on the same chromosome, it was manually inspected to check if it was truly absent in the other genome. Merged ORFs and misannotations were not considered in further analysis. These alignments were also used to identify copies and homologues of the genes identified as truly absent in the other genome.

Gene ontology analysis was performed using the Gene Ontology term finder of SGD using the list of unique genes as the query set and all annotated genes as the background set of genes for each genome (Additional file 2A and 2C). The ORFs of genes identified as present in S288C but absent in CEN.PK113-7D in previously made lists (Daran-Lapujade *et al.* 2003, Nijkamp *et al.* 2012) were obtained from SGD. The ORFs were aligned both ways to ORFs from SGD identified as unique to S288C in this study using BLASTN. Genes with alignments of at least half the query length and with a sequence identity of at least 95% were interpreted as confirmed by the other data set. In order to analyze the origin of genes identified as unique to S288C, these ORFs were aligned using BLASTN to 481 genome assemblies of various *S. cerevisiae* strains obtained from NCBI (Additional file 3) and alignments of at least 50% of the query were considered. The top alignments were selected based on the highest sequence ID and only one alignment per strain was counted per gene.

### Chromosome translocation analysis

Reads supporting the original and translocated genomic architectures of chromosomes III and VIII were identified via read alignment of raw nanopore reads. First, the translocation breakpoints coordinates were calculated based on whole-genome alignment of CEN.PK113-7D Delft assembly to S288C with MUMmer. A modified version of S288C was created containing the normal architectures of all 16 chromosomes and the mitochondrial genome plus the translocated architecture of chromosomes III-VIII and VIII-III. The first nearest unique flanking genes at each breakpoint were determined using BLASTN (version 2.2.31+) (English *et al.* 2012, Zhang *et al.* 2000) in reference to both S288C and the Delft CEN.PK113-7D nanopore assembly. Raw nanopore reads from CEN.PK113-7D Delft and Frankfurt were aligned to the modified version of S288C and nanopore reads that spanned the translocation breakpoints as well as the unique flanking sequences were extracted. Supporting reads were validated by re-aligning them to the modified version of S288C using BLASTN.

## Results

### Sequencing on a single nanopore flow cell enables near-complete genome assembly

To obtain a complete chromosome level *de novo* assembly of *Saccharomyces cerevisiae* CENPK113-7D, we performed long read sequencing on the Oxford Nanopore Technology’s (ONT) MinION platform. A fresh sample of CEN.PK113-7D was obtained from the original distributer Dr. P. Kötter (further referred to as “CEN.PK113-7D Frankfurt”), cultured in a single batch on YPD medium and genomic DNA was extracted. CEN.PK113-7D Frankfurt was sequenced on a single R9 (FLO-MIN104) chemistry flow cell using the 2D ligation kit for the DNA libraries producing more than 49x coverage of the genome with an average read-length distribution of 10.0 Kbp (Supplementary Figure S1) and an estimated error rate of 10% (Supplementary Figure S2). We used Canu (Koren *et al.* 2017) to produce high-quality de novo assemblies using only nanopore data. Before correcting for misassemblies, the assembly contained a total of 21 contigs with an N50 of 756 Kbp (Supplementary Table S1). This represented a 19-fold reduction in the number of contigs and a 15-fold increase of the N50 in comparison to the short-read-only assembly of the first CEN.PK113-7D draft genome version (Nijkamp *et al.* 2012) (Table 1).

**Table 1.**
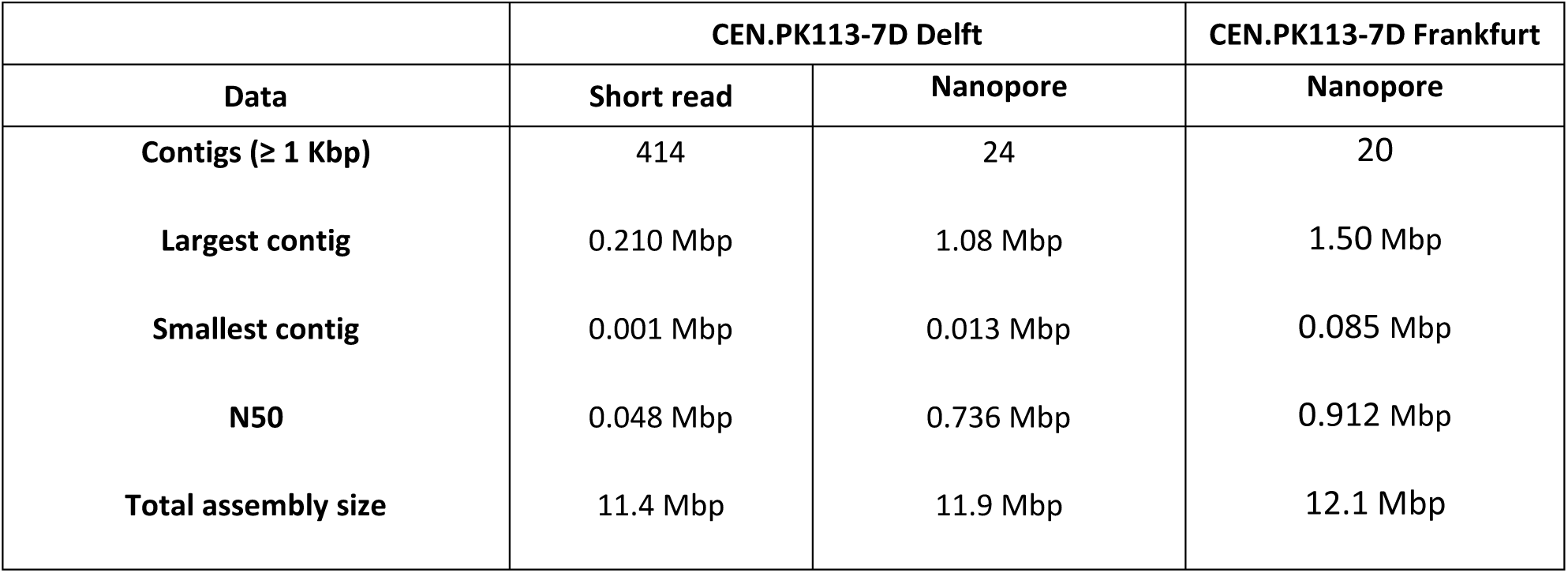
Comparison of 454/Illumina and nanopore *de novo* assemblies of CEN.PK113-7D. Summary of *de novo* assembly metrics of CEN.PK113-7D Delft and CEN.PK113-7D Frankfurt. For the short-read assembly, only contigs of at least 1 Kbp are shown (Nijkamp *et al.* 2012). The nanopore assembly of CEN.PK113-7D Delft is uncorrected for misassemblies while CEN.PK113-7D Frankfurt was corrected for misassemblies.

|  | CEN.PK113-7D Delft |  | CEN.PK113-7D Frankfurt |
| --- | --- | --- | --- |
| Data | Short read | Nanopore | Nanopore |
| Contigs ($\geq 1$ Kbp) | 414 | 24 | 20 |
| Largest contig | 0.210 Mbp | 1.08 Mbp | 1.50 Mbp |
| Smallest contig | 0.001 Mbp | 0.013 Mbp | 0.085 Mbp |
| N50 | 0.048 Mbp | 0.736 Mbp | 0.912 Mbp |
| Total assembly size | 11.4 Mbp | 11.9 Mbp | 12.1 Mbp |

Most chromosomes of the nanopore *de novo* assembly are single contigs and are flanked by telomere caps. Genome completeness was determined by alignment to the manually-curated reference genome of the strain S288C (version R64, Genbank ID: 285798) (Supplementary Table S2). The two largest yeast chromosomes, IV and XII, were each split in two separate contigs, and two additional contigs (31 and 38 Kbp in length) corresponded to unplaced subtelomeric fragments. In particular, the assembly for chromosome XII was interupted in the *RDN1* locus—a repetitive region consisting of gene encoding ribosomal RNA estimated to be more than 1-Mbp long (Venema and Tollervey 1999). Since no reads were long enough to span this region, the contigs were joined with a gap.

Manual curation resolved chromosome III, chromosome IV and the mitochondrial genome. Chromosome IV was fragmented into two contigs at locus of 11.5 Kbp containing two Ty-elements in S288C (coordinates 981171-992642). Interestingly, the end of the first contig and the start of the second contig have 8.8 Kbp of overlap (corresponding to the two Ty-elements) and one read spans the repetitive Ty-elements and aligns to unique genes on the left and right flanks (*EXG2* and *DIN7*, respectively). We therefore joined the contigs without missing sequence resulting in a complete assembly of chromosome IV. For chromosome III, the last ∼27 Kbp contained multiple telomeric caps next to each other. The last ∼10 Kbp had little to no coverage when re-aligning raw nanopore reads to the assembly (Supplementary Figure S3). The coordinates for the first telomeric cap were identified and the remaining sequence downstream was removed resulting in a final contig of size of 347 Kbp. The original contig corresponding to the mitochondrial genome had a size of 104 Kbp and contained a nearly identical ∼20 Kbp overlap corresponding to start of the *S. cerevisiae* mitochondrial genome (i.e. origin of replication) (Supplementary Figure S4). This is a common artifact as assembly algorithms generally have difficulties reconstructing and closing circular genomes (McIlwain *et al.* 2016, Venema and Tollervey 1999). The coordinates of the overlaps were determined with Nucmer (Kurtz *et al.* 2004) and manually joined resulting to a final size of 86,616 bp.

Overall, the final CEN.PK113-7D Frankfurt assembly contained 15 chromosome contigs, 1 chromosome scaffold, the complete mitochondrial contig, the complete 2-micron plasmid and two unplaced telomeric fragments, adding up to a total of 12.1 Mbp (Table 1 and Supplementary Table S3). Of the 16 chromosomes, 11 were assembled up until both telomeric caps, four were missing one of the telomere caps and only chromosome X was missing both telomere caps. Based on homology with S288C, the missing sequence was estimated not to exceed 12 kbp for each missing (sub)telomeric region. Furthermore, we found a total of 46 retrotransposons Ty-elements: 44 were from the *Pseudoviridae* group (30 *Ty1*, 12 *Ty2*, 1 *Ty4*, and 1 *Ty5*) and 2 from *Metaviridae* group (*Ty3*). The annotated nanopore assembly of CEN.PK113-7D Frankfurt is available at NCBI under the bioproject accession number PRJNA393501.

### Comparison of the nanopore and short-read assemblies of CEN.PK113-7D

We compared the nanopore assembly of CEN.PK113-7D to a previously published version to quantify the improvements over the current state-of-the art (Nijkamp *et al.* 2012). Alignment of the contigs of the short-read assembly to the nanopore assembly revealed 770 Kbp of previously unassembled sequence, including the previously unassembled mitochondrial genome (Additional file 4A). This gained sequence is relatively spread out over the genome (Figures 1A and 1B) and contained as much as 284 chromosomal gene annotations (Additional file 4B). Interestingly, 69 out of 284 genes had paralogs, corresponding to a fraction almost twice as high as the 13% found in the whole genome of S288C (Wolfe and Shields 1997). Gene ontology analysis revealed an enrichment in the biological process of cell aggregation (9.30x10^−4^); in the molecular functions of mannose binding (P=3.90x10^−4^) and glucosidase activity (P=7.49x10^−3^); and in the cellular components of the cell wall (P=3.41x10^−7^) and the cell periphery component (P=5.81x10^−5^). Some newly-assembled genes are involved in central carbon metabolism, such as *PDC5*. In addition, many of the added genes are known to be relevant in industrial applications including hexose transporters such as *HXT* genes and sugar polymer hydrolases such as *IMA* and *MALx2* genes; several genes relevant for cellular metal homeostasis, such as *CUP1-2* (linked to copper ion tolerance) and *FIT1* (linked to iron ion retention); genes relevant for nitrogen metabolism in medium rich or poor in specific amino acids, including amino acid transporters such as *VBA5*, amino acid catabolism genes such as *ASP3-4* and *LEU2* and amino-acid limitation response genes such as many *PAU* genes; several *FLO* genes which are responsible for calcium-dependent flocculation; and various genes linked to different environmental stress responses, such as *HSP* genes increasing heat shock tolerance and *RIM101* increasing tolerance to high pH.

**Figure 1:**
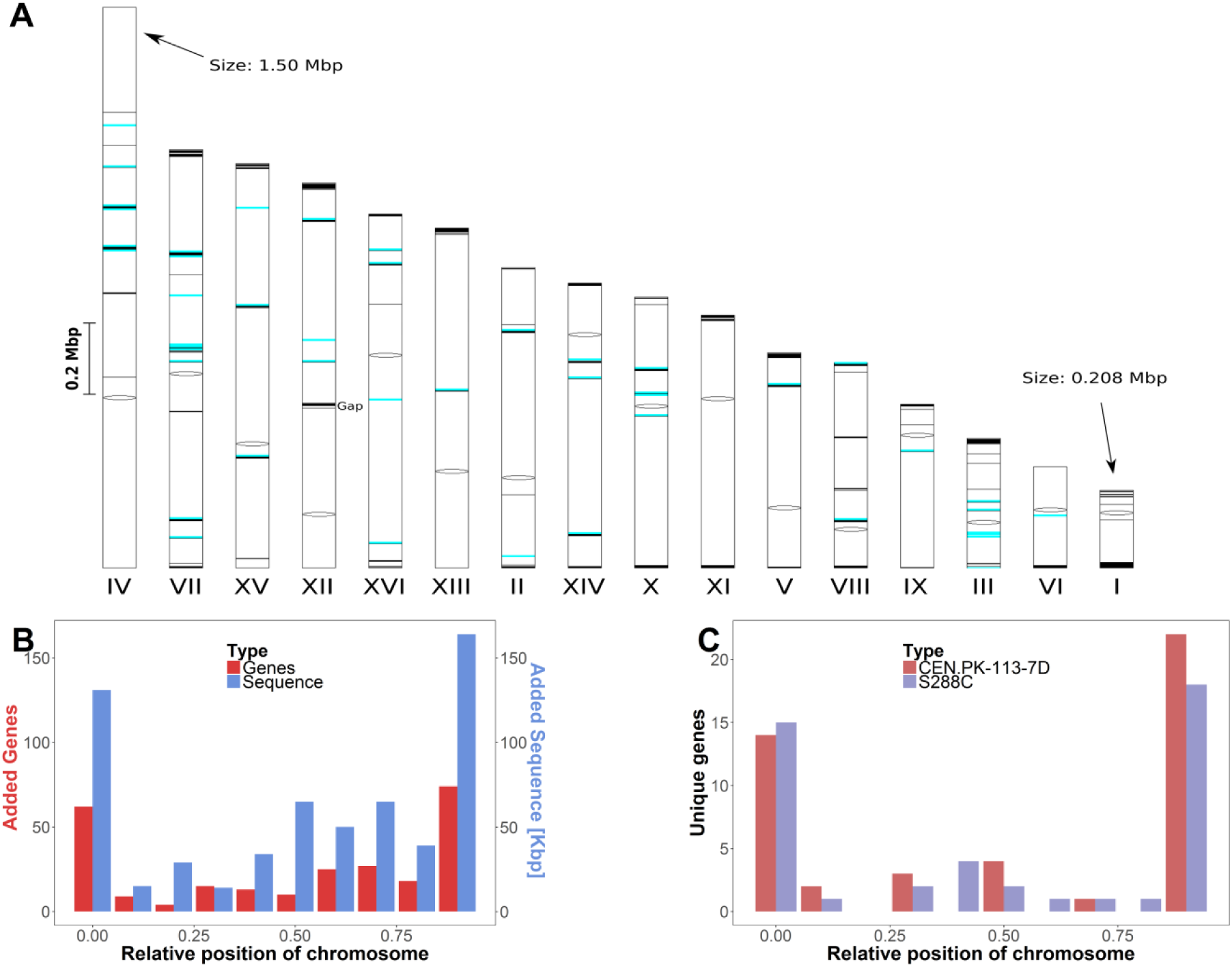
Overview of gained and lost sequence and genes in the CEN.PK113-7D Frankfurt nanopore assembly relative to the short-read CEN.PK113-7D assembly and to the genome of S288C. The two unplaced subtelomeric contigs and the mitochondrial DNA were not included in this figure. **(1A) Chromosomal location of sequence assembled in the nanopore assembly which was not assembled using short-read data**. The sixteen chromosome contigs of the nanopore assembly are shown. Chromosome XII has a gap at the *RDN1* locus, a region estimated to contain more than 1 Mbp worth of repetitive sequence (Venema and Tollervey 1999). Centromeres are indicated by black ovals, gained sequence relative to the short-read assembly is indicated by black marks and 46 identified retrotransposon Ty-elements are indicated by blue marks. The size of all chromosomes and marks is proportional to their corresponding sequence size. In total 611 Kbp of sequence was added within the chromosomal contigs. **(1B) Relative chromosome position of sequences and genes assembled on chromosome contigs of the nanopore assembly which were not assembled using short-read data**. The positions of added sequence and genes were normalized to the total chromosome size. The number of genes (red) and the amount of sequence (cyan) over all chromosomes are shown per tenth of the relative chromosome size. **(1C) Relative chromosome position of gene presence differences between S288C and CEN.PK113-7D.** The positions of the 45 genes identified as unique to CEN.PK113-7D and of the 44 genes identified as unique to S288C were normalized to the total chromosome size. The number of genes unique to CEN.PK113-7D (red) and to S288C (purple) are shown per tenth of the relative chromosome position.

To evaluate whether some previously assembled sequence was missing in the nanopore assembly, we aligned the nanopore contigs to the short-read assembly (Nijkamp *et al.* 2012). Less than 6 Kbp of sequence of the short-read assembly was not present in the nanopore assembly, distributed over 13 contigs (Additional file 4C). Only two ORFs were missing: the genes *BIO1* and *BIO6* (Additional file 4D). Alignment of *BIO1* and *BIO6* sequences to the nanopore assembly showed that the right-end of the chromosome I contig contains the first ∼500 nt of *BIO1*. While *BIO1* and *BIO6* were present in the nanopore sequences, they are absent in the final assembly likely due to the lack of long-enough reads to resolve the repetitive nature of this subtelomeric region.

Overall an additional 770 Kbp sequence containing 284 genes was gained, while 6 Kbp containing two genes was not captured compared to the previous assembly. In addition, the reduction from over 700 to only 20 contigs clearly shows that the nanopore assembly is much less fragmented than the short-read assembly (Table 1).

### Comparison of the Nanopore assembly of CEN.PK113-7D to S288C

To identify unique and shared genes between CEN.PK113-7D and S288C, we compared annotations made using the same method for both genomes (Additional Files 2A and 2C). We identified a total of 45 genes unique to CEN.PK113-7D and 44 genes unique to S288C (Additional Files 2B and 2D). Genes located in regions that had no assembled counterpart in the other genome were excluded; 20 for S288C and 27 for CEN.PK113-7D. Interestingly, the genes unique to either strain and genes present on different chromosomes were found mostly in the outer 10% of the chromosomes, indicating that the subtelomeric regions harbor most of the genetic differences between CEN.PK113-7D and S288C (Figure 1C).

In order to validate the genes identified as unique to S288C, we compared them to genes identified as absent in CEN.PK113-7D in previous studies (Additional file 2D, Table 2). 25 genes of S288C were identified as absent in CEN.PK113-7D by array comparative genomic hybridization (aCGH) analysis (Daran-Lapujade *et al.* 2003) and 21 genes were identified as absent in CEN.PK113-7D based on short-read WGS (Nijkamp *et al.* 2012). Of these genes, 19 and 10 respectively were identified as genes in S288C by our annotation pipeline and could be compared to the genes we identified as unique to S288C. While 19 of these 29 genes were also absent in the nanopore assembly, the remaining 10 genes were fully assembled and annotated, indicating they were erroneously identified as missing (Table 2).

**Table 2:** Presence in the nanopore assembly of genes identified as absent in CEN.PK113-7D in previous research. For genes identified as absent in CEN.PK113-7D in two previous studies, the absence or presence in the nanopore assembly of CEN.PK113-7D is shown. 25 genes were identified previously by array comparative genome hybridisation (Daran-Lapujade *et al.* 2003) and 21 genes were identified by short-read genome assembly (Nijkamp *et al.* 2012). Genes which were not annotated by MAKER2 in S288C could not be analysed. Genes with an alignment to genes identified as missing in the nanopore assembly of at least 50% of the query length and 95% sequence identity were confirmed as being absent, while those without such an alignment were identified as present. The presence of these genes was verified manually, which revealed the misanotation of YPL277C as YOR389W.

|  | Not analysed | Absent in assembly | Present in assembly |
| --- | --- | --- | --- |
| Daran-Lapujade et al | YAL064C-A, YAL066W, YAR047C, YHL046W-A, YIL058W, YOL013W-A | YAL065C, YAL067C, YBR093C, YCR018C, YCR105W, YCR106W, YDR038C, YDR039C, YHL047C, YHL048W, YNR070W, YNR071C and YNR074C | YAL069W, YDR036C, YDR037W, YJL165C, YNR004W, and YPL277C (misannotated as YOR389W) |
| Nijkamp et al | Q0140, YDR543C, YDR544C, YDR545W, YIL046W-A, YLR154C-H, YLR156C-A, YLR157C-C, YLR159C-A, YOR029W and YOR082C | YBR093C, YCR040W, YCR041W, YDR038C, YDR039C and YDR040C | YDR036C, YHL008C, YHR056C and YLR055C |

In order to determine if the genes unique to S288C have homologues elsewhere in the genome of CEN.PK113-7D or if they are truly unique, we aligned the ORFs of the 44 genes identified as unique in S288C to the ORFs in the naopore CEN.PK113-7D assembly. 26 genes were completely absent in the CEN.PK113-7D assembly, while the remaining 18 genes aligned to between 1 and 20 ORFs each in the genome of CEN.PK113-7D with more than 95% sequence identity, indicating they may have close homologues or additional copies in S288C (Additional file 2D). Gene ontology analysis revealed no enrichment in biological process, molecular functions or cell components of the 26 genes without homologues in CEN.PK113-7D. Five genes without homologues were labelled as putative. However, there were many genes encoding proteins relevant for fitness under specific industrial conditions, such as *PHO5* which is part of the response to phosphate scarcity, *COS3* linked to salt tolerance, *ADH7* linked to acetaldehyde tolerance, *RDS1* linked to resistance to cycloheximide, *PDR18* linked to ethanol tolerance and *HXT17* which is involved in hexose sugar uptake (Additional file 2D). In addition, we confirmed the complete absence of *ENA2* and *ENA5* in CEN.PK113-7D which are responsible for lithium sensitivity of CEN.PK113-7D (Daran-Lapujade *et al.* 2009).

Conversely, to determine if the genes unique to CEN.PK113-7D have homologues elsewhere in the genome of S288C or if they are truly unique, we aligned the ORFs of the 45 genes identified as unique in CEN.PK113-7D to the ORFs of S288C. A set of 16 genes were completely absent in S288C, while the remaining 29 aligned to between one and 16 ORFs each in the genome of S288C with more than 95% sequence (Additional File 2D). Gene ontology analysis revealed no enrichment in biological processes, molecular functions or cell components of the 16 genes unique to CEN.PK113-7D without homologues. However, among the genes without homologues a total of 13 were labelled as putative. The presence of an additional copy of *IMA1*, *MAL31* and *MAL32* on chromosome III was in line with the presence of the *MAL2* locus which was absent in S288C. Interestingly the sequence of *MAL13,* which belongs to this locus, was divergent enough from other *MAL*-gene activators to not be identified as homologue. Additionally, when performing the same analysis on the 27 genes on the two unplaced contigs of the CEN.PK113-7D assembly, 7 of them did not align to any gene of S288C with more than 95% sequence identity, indicating these unplaced telomeric regions are highly unique to CEN.PK113-7D.

Since the genome of CEN.PK113-7D contains 45 ORFs which are absent in S288C, we investigated their origin by aligning them against all available *S. cerevisiae* nucleotide data at NCBI (Additional File 3). For each ORF, we report the strains to which they align with the highest sequence identity and the sequence identity relative to S288C in Additional File 2B. For most genes, several strains aligned equally well with the same sequence identity. For 13 ORFs S288C is among the best matches, indicating these ORFs may come from duplications in the S288C genome. However, S288C is not among the best matches for 32 ORFs. In these, laboratory strain “SK1” is among the best matches 9 times, the west African wine isolate “DBVPG6044”appears 8 times, laboratory strain “W303” appears 7 times, the Belgian beer strain “beer080” appears 3 times and the Brazilian bioethanol strain “bioethanol005”appears 3 times. Interestingly, some grouped unique genes are most related to specific strains. For example, the unique genes identified on the left subtelomeric regions of chromosome XVI (YBL109W, YHR216W and YOR392) and of chromosome VIII (YJL225C and YOL161W) exhibited the highest similarity to DBVPG6044. Similarly, the right end of the subtelomeric region of chromosome III (YPL283W-A and YPR202) and of chromosome XI ((YPL283W-A and YLR466W) were most closely related to W303.

Interestingly, the nanopore assembly revealed a duplication of *LEU2*, a gene involved in synthesis of leucine which can be used as an auxotrophy marker. In the complete reference genome of *S. cerevisiae* S288C, both *LEU2* and *NFS1* are unique, neighboring genes located chromosome III. However, gene annotations of the assemblies and raw nanopore reads support additional copies of *LEU2* and *NFS1* in CEN.PK113-7D located on chromosome VII (Figure 2). The additional copy contained the complete *LEU2* sequence but only ∼0.5 kb of the 5’ end of *NFS1*. In CEN.PK113-7D and S288C, the *LEU2* and *NFS1* loci in chromosome III were located adjacent to Ty-elements. Two such Ty-elements were also found flanking the additional *LEU2* and *NFS1* loci in chromosome VII (Figure 2). It is likely that the duplication was the result of a translocation based on homology of the Ty-elements which resulted in local copy number increase during its strain development program (Entian and Kötter 2007).

**Figure 2:**
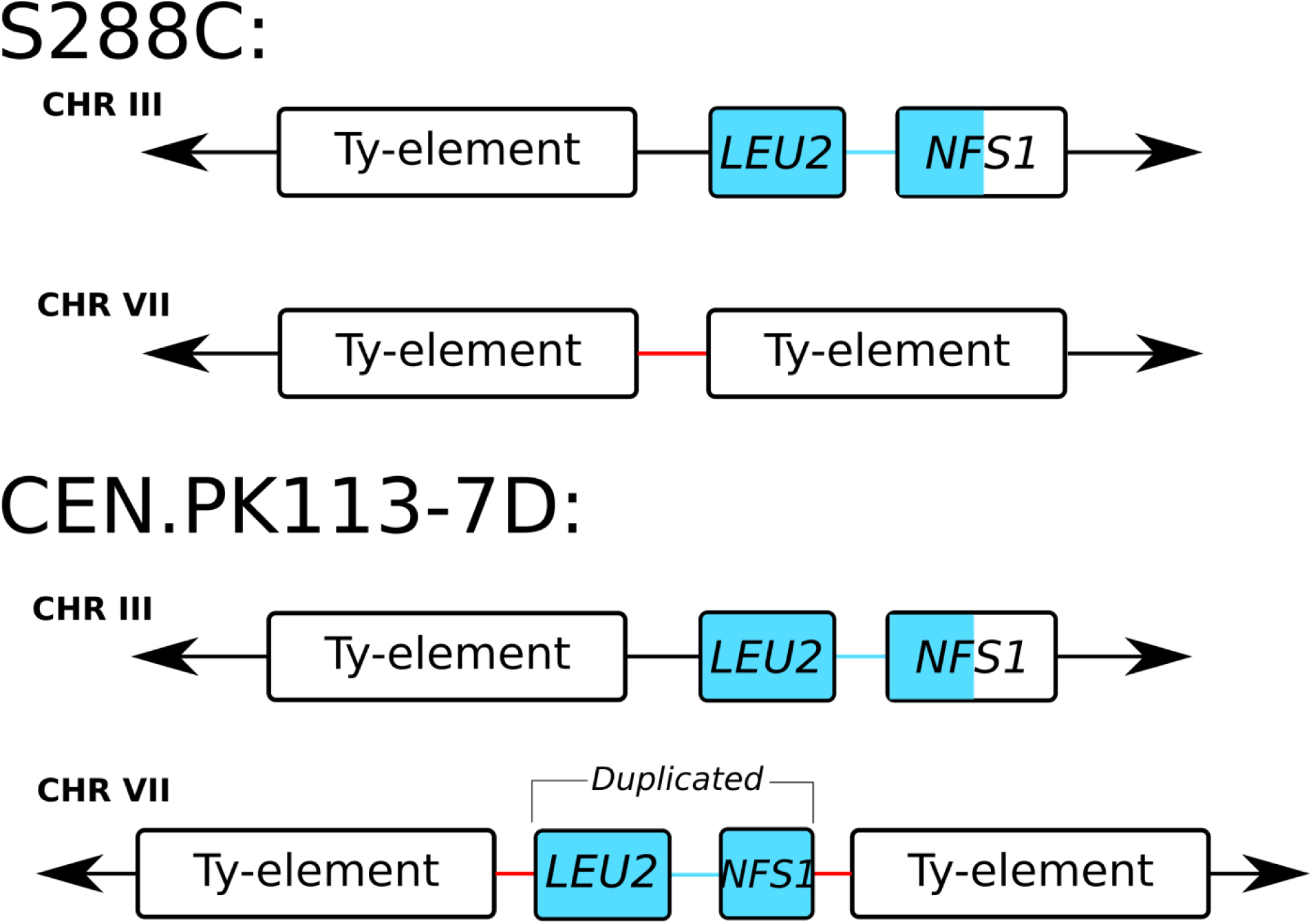
LEU2 and NFS1 duplication in chromosome VII of CEN.PK113-7D. The nanopore assembly contains a duplication of *LEU2* and part of *NFS1* in CEN.PK113-7D. In S288C, the two genes are located in chromosome III next to a Ty element. In CEN.PK113-7D, the two genes are present in chromosome III and in chromosome VII. The duplication appears to be mediated by Ty-elements. Note that the additional copy in chromosome VII is present in between two Ty-elements and contains only the first ∼500 bp of *NFS1*. The duplication is supported by long-read data that span across the *LEU2*, *NFS1*, the two Ty-elements, and the neighboring flanking genes (not shown).

### Long-read sequencing data reveals chromosome structure heterogeneity in CEN.PK113-7D Delft

CEN.PK113-7D has three confirmed *MAL* loci encoding genes for the uptake and hydrolysis of maltose: *MAL1* on chromosome VIII, *MAL2* on chromosome III and *MAL3* on chromosome II (Additional file 2A). A fourth *MAL* locus was identified in previous research on chromosome XI based on contour-clamped homogeneous electric field electrophoresis (CHEF) and southern blotting with a probe for *MAL* loci (Nijkamp *et al.* 2012). However, the nanopore assembly revealed no additional *MAL* locus despite the complete assembly of Chromosome XI. The CEN.PK113-7D stock in which the fourth *MAL* locus was obtained from Dr P. Kötter in 2001 and stored at −80°C since (further referred to as “CEN.PK113-7D Delft”). In order to investigate the presence of the potential *MAL* locus, we sequenced CEN.PK113-7D Delft using nanopore MinION sequencing. Two R7.3 flow cells (FLO-MIN103) produced 55x coverage with an average read-length distribution of 8.5 Kbp and an R9 flow cell (FLO-MIN103) produced 47x coverage with an average read-length distribution of 3.2 Kbp (Supplementary Figure S1). The error rate was estimated to be 13% (Supplementary Figure S4) after aligning the raw nanopore reads to the CEN.PK113-7D Frankfurt assembly. These reads were assembled into 24 contigs with an N50 of 736 Kbp (Supplementary Table S1).

Alignment of the assembly of CEN.PK113-7D Delft to the Frankfurt assembly showed evidence of a translocation between chromosomes III and VIII (Supplementary Figure S5). The assembly thus suggested the presence of two new chromosomes: chromosomes III-VIII of size 680 Kbp and chromosome VIII-III of size 217 Kbp (Figure 3). The translocation occurred between Ty-element YCLWTy2-1 on chromosome III and long terminal repeats YHRCdelta5-7 on chromosome VIII. These repetitive regions are flanked by unique genes *KCC4* and *NFS1* on chromosome III and *SPO13* and *MIP6* on chromosome VIII (Figure 3). Nanopore reads spanning the whole translocated or non-translocated sequence anchored in the unique genes flanking them were extracted for CEN.PK113-7D Delft and Frankfurt. A total of eight reads from CEN.PK113-7D Delft supported the translocated chromosome III-VIII architecture (largest read was 39 Kbp) and one 19 Kbp read supported the normal chromosome III architecture. For CEN.PK113-7D Frankfurt, we found only one read of size 23 Kbp that supported the normal chromosome III architecture but we found no reads that supported the translocated architectures. This data suggested that CEN.PK113-7D Delft is in fact a heterogeneous population containing cells with recombined chromosomes III and VIII and cells with original chromosomes III an VIII. As a result, in addition to the *MAL2* locus on chromosome III, CEN.PK113-7D Delft harboured a *MAL2* locus on recombined chromosome III-VIII. As the size of recombined chromosome III-VIII was close to chromosome XI, the *MAL2* locus on chromosome III-VIII led to misidentification of a *MAL4* locus on chromosome XI (Nijkamp *et al.* 2012). By repeating the CHEF gel and southern blotting for MAL loci on several CEN.PK113-7D stocks, the *MAL2* on the translocated chromosomes III-VIII was shown to be present only in CEN.PK113-7D Delft, demonstrating that there was indeed chromosome structure heterogeneity (Additional File 5).

**Figure 3:**
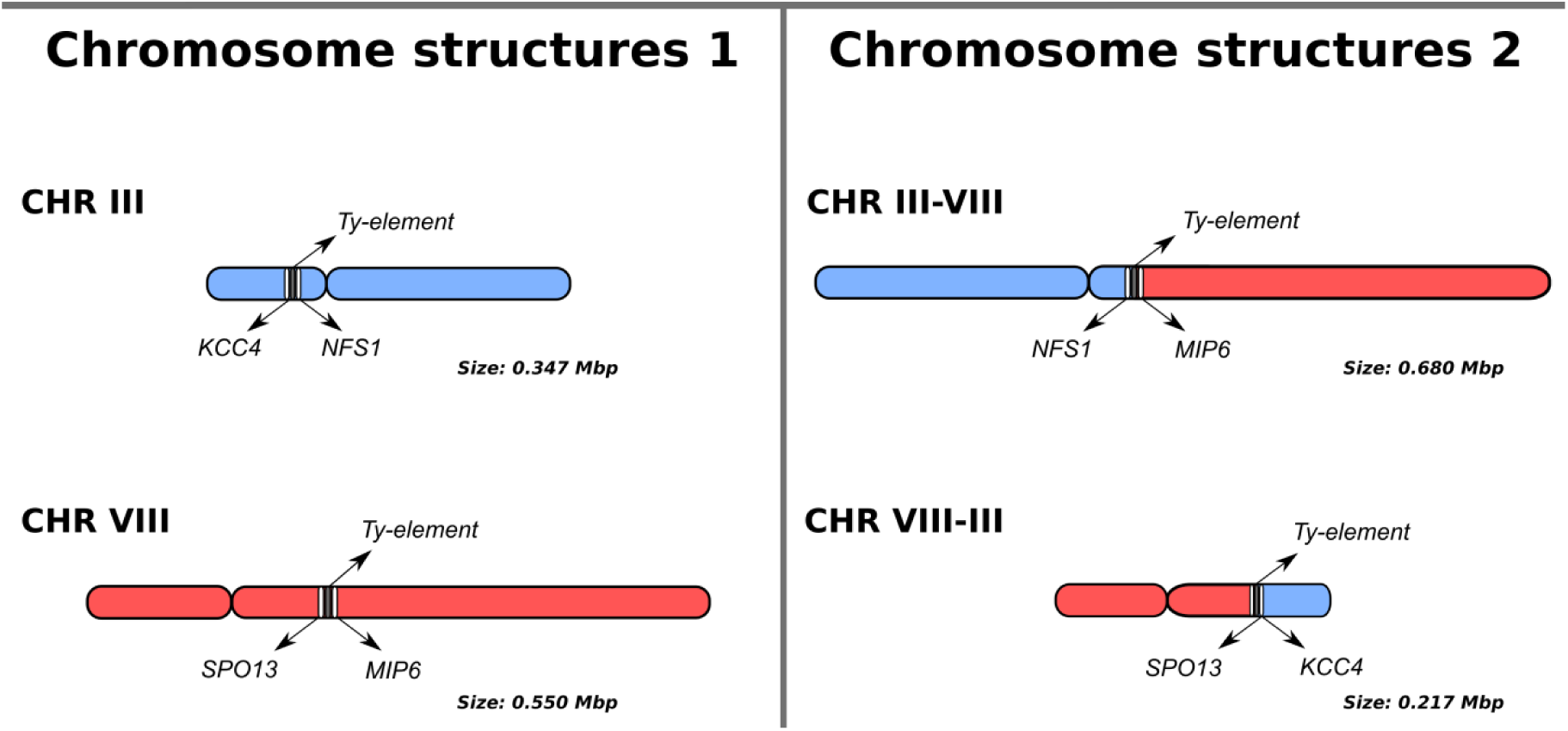
Overview of chromosome structure heterogeneity in CEN.PK113-7D Delft for CHRIII and CHRVIII which led to the misidentification of a fourth MAL locus in a previous short-read assembly study of the genome of CEN.PK113-7D. Nanopore reads support the presence of two chromosome architectures: the normal chromosomes III and VIII (left panel) and translocated chromosomes III-VIII and VIII-III (right panel). The translocation occurred in Ty-elements, large repetitive sequences known to mediate chromosomal translocations in *Saccharomyces* species (Fischer *et al.* 2000). Long-reads are required to diagnose the chromosome architecture via sequencing: the repetitive region between *KCC4* to *NFS1* in chromosome III exceeds 15 Kbp, while the region between *SPO13* and *MIP6* in chromosome VIII is only 1.4 Kbp long. For the translocated architecture, the region from *NFS1* to *MIP6* in chromosome III-VIII exceeds 16 Kbp and the distance from *SPO13* to *KCC4* in chromosome VIII-III is nearly 10 Kbp.

## Discussion

In this study, we obtained a near-complete genome assembly of *S. cerevisiae* strain CEN.PK113-7D using only a single R9 flow cell on ONT’s MinION sequencing platform. 15 of the 16 chromosomes as well as the mitochondrial genome and the 2-micron plasmid were assembled in single, mostly telomere-to-telomere, contigs. This genome assembly is remarkably unfragmented, even when compared with other *S. cerevisiae* assemblies made with several nanopore technology flow cells, in which 18 to 105 chromosomal contigs were obtained (Istace *et al.* 2017, McIlwain *et al.* 2016). Despite the long read lengths obtained by Nanopore sequencing, the ribosomal DNA locus in chromosome XII could not be completely resolved. In practice, this would require reads exceeding 1 Mb in length, which current technology cannot yet deliver.

The obtained nanopore assembly is of vastly superior quality to the previous short-read-only assembly of CEN.PK113-7D that was fragmented into over 700 contigs (Nijkamp *et al.* 2012). In addition to the lesser fragmentation, the addition of 770 Kbp of previously unassembled sequence led to the identification and accurate placement of 284 additional ORFs spread out over the genome. These newly assembled genes showed overrepresentation for cell wall and cell periphery compartmentalization and relate to functions such as sugar utilization, amino acid uptake, metal ion metabolism, flocculation and tolerance to various stresses. While many of these genes are already present in the short-read assembly of CEN.PK113-7D, copy number was shown to be an important factor determining the adaptation of strains to specific growth conditions (Brown *et al.* 2010). The added genes may therefore be very relevant for the specific physiology of CEN.PK113-7D under different industrial conditions (Brown *et al.* 2010). The ability of nanopore sequencing to distinguish genes with various similar copies is crucial in *S. cerevisiae* as homologues are frequent particularly in subtelomeric regions, and paralogues are widespread due to a whole genome duplication in its evolutionary history (Wolfe and Shields 1997). Besides the added sequence, 6 Kbp of sequence of the short-read assembly was not present in the nanopore assembly, mostly consisting of small unplaced contigs. Notably the absence of *BIO1* and *BIO6* in the assembly was unexpected, as it constituted a marked difference between CEN.PK113-7D and many other strains which enables biotin prototrophy (Bracher *et al.* 2017). Both genes were present in the nanopore reads, but were unassembled likely due to the lack of reads long-enough to resolve this subtelomeric region (a fragment of *BIO1* is located at the right-end of chromosome I). Targeted long-read sequencing in known gaps of a draft assembly followed by manual curation could provide an interesting tool to obtain complete genome assemblies (Loose *et al.* 2016). Alternatively, a more complete assembly could be obtained by maximizing read length. The importance of read length is illustrated by the higher fragmentation of the CEN.PK113-7D Delft assembly compared to the Frankfurt one, which was based on reads with lower length distribution despite higher coverage and similar error rate (Table 1, Supplementary Figures S1 and S5). Read-length distribution in nanopore sequencing is highly influenced by the DNA extraction method and library preparation (Supplementary Figure S1). The mitochondrial genome was completely assembled, which is not always possible with nanopore sequencing (Giordano *et al.* 2017, Istace *et al.* 2017, McIlwain *et al.* 2016). Even with identical DNA extraction and assembly methods, the mitochondrial genome cannot always be assembled, as illustrated by its absence in the assembly of CEN.PK113-7D Delft. Overall, the gained sequence in the nanopore assembly far outweighs the lost sequence relative to the previous assembly, and the reduction in number of contigs presents an important advantage.

The use of long read sequencing enabled the discovery of a translocation between chromosomes III and VIII, which led to the misidentification of a fourth MAL locus on chromosome XI of CEN.PK113-7D (Nijkamp *et al.* 2012). Identification of this translocation required reads to span at least 12 Kbp due to the large repetitive elements surrounding the translocation breakpoints, explaining why it was previously undetected. While the translocation did not disrupt any coding sequence and is unlikely to cause phenotypical changes (Naseeb *et al.* 2016), there may be decreased spore viability upon mating with other CEN.PK strains. Our ability to detect structural heterogeneity within a culture shows that nanopore sequencing could also be valuable in detecting structural variation within a genome between different chromosome copies, which occurs frequently in aneuploid yeast genomes (Gorter de Vries *et al.* 2017). These results highlight the importance of minimal propagation of laboratory microorganisms to warrant genome stability and avoid heterogeneity which could at worst have an impact on phenotype and interpretation of experimental results.

The nanopore assembly of CEN.PK113-7D constitutes a vast improvement of its reference genome which should facilitate its use as a model organism. The elucidation of various homologue and paralogue genes is particularly relevant as CEN.PK113-7D is commonly used as a model for industrial *S. cerevisiae* applications for which gene copy number frequently plays an important role (Brown *et al.* 2010, Gorter de Vries *et al.* 2017). Using the nanopore assembly as a reference for short-read sequencing of strains derived from CEN.PK113-7D will yield more complete and more accurate lists of SNPs and other mutations, facilitating the identification of causal mutations in laboratory evolution or mutagenesis experiments. Therefore, the new assembly should accelerate elucidation of the genetic basis underlying the fitness of *S. cerevisiae* in various environmental conditions, as well as the discovery of new strain improvement strategies for industrial applications (Oud *et al.* 2012).

## Acknowledgements

The authors would like to thank Dr. P. Kötter for sending CEN.PK113-7D Frankfurt, Dr. Kirsten Benjamin for sending CEN.PK113-7D Amyris and Dr. Verena Siewers for sending CEN.PK113-7D Chalmers. We are thankful to Prof. Jack T. Pronk (Delft University of Technology) and Dr. Niels Kuijpers (HEINEKEN Supply Chain B.V.) for their critical reading of the manuscript.

This work was performed within the BE-Basic R&D Program (http://www.be-basic.org/), which was granted an FES subsidy from the Dutch Ministry of Economic Affairs, Agriculture and Innovation (EL&I). Anja Brickwedde was funded by the Seventh Framework Programme of the European Union in the frame of the SP3 people support for training and career development of researchers (Marie Curie), Networks for Initial Training (PITN-GA-2013 ITN-2013-606795) YeastCell.

## Author’s contribution

PdlTC and ARGdV extracted high molecular weight DNA for Illumina and MinION sequencing. PdlTC performed Illumina sequencing. AS and MW constructed MinION sequencing libraries and performed MinION genome sequencing. ARGdV and AB performed the CHEF and Southern-blot hybridization. AS, ARGdV and MvdB performed the bioinformatics analysis. AS, ARGdV, MvdB, JMGD and TA were involved in the data analysis and AS, ARGdV, JMGD and TA wrote the manuscript. JMGD and TA supervised the study. All authors read and approved the final manuscript.

## References

Bergström A, Simpson JT, Salinas F et al. (2014) A high-definition view of functional genetic variation from natural yeast genomes. Mol Biol Evol 31: 872–88.

Bracher JM, de Hulster E, Koster CC et al. (2017) Laboratory evolution of a biotin-requiring *Saccharomyces cerevisiae* strain for full biotin prototrophy and identification of causal mutations. Appl Environ Microbiol AEM. 00892-17.

Brown CA, Murray AW, Verstrepen KJ (2010) Rapid expansion and functional divergence of subtelomeric gene families in yeasts. Current biology: CB 20: 895–903.

Camacho C, Coulouris G, Avagyan V et al. (2009) BLAST+: architecture and applications. BMC bioinformatics 10: 421.

Canelas AB, Harrison N, Fazio A et al. (2010) Integrated multilaboratory systems biology reveals differences in protein metabolism between two reference yeast strains. Nat Commun 1: 145.

Carlson M, Celenza JL, Eng FJ (1985) Evolution of the dispersed *SUC* gene family of *Saccharomyces* by rearrangements of chromosome telomeres. Mol Cell Biol 5: 2894–902.

Cherry JM, Hong EL, Amundsen C et al. (2012) *Saccharomyces* Genome Database: the genomics resource of budding yeast. Nucleic Acids Res 40: D700–5.

Daran-Lapujade P, Daran J-MG, Kötter P et al. (2003) Comparative genotyping of the *Saccharomyces cerevisiae* laboratory strains S288C and CEN. PK113-7D using oligonucleotide microarrays. Fems Yeast Res 4: 259–69.

Daran-Lapujade P, Daran J-MG, Luttik MAH et al. (2009) An atypical *PMR2* locus is responsible for hypersensitivity to sodium and lithium cations in the laboratory strain *Saccharomyces cerevisiae* CEN. PK113-7D. Fems Yeast Res 9: 789–92.

Denayrolles M, de Villechenon EP, Lonvaud-Funel A et al. (1997) Incidence of *SUC-RTM* telomeric repeated genes in brewing and wild wine strains of *Saccharomyces*. Curr Genet 31: 457–61.

English AC, Richards S, Han Y et al. (2012) Mind the gap: upgrading genomes with Pacific Biosciences RS long-read sequencing technology. PLoS One 7: e47768.

Entian K-D, Kötter P (2007) 25 Yeast genetic strain and plasmid collections. Method Microbiol 36: 629–66.

Fischer G, James SA, Roberts IN et al. (2000) Chromosomal evolution in *Saccharomyces*. Nature 405: 451–4.

Giordano F, Aigrain L, Quail MA et al. (2017) *De novo* yeast genome assemblies from MinION, PacBio and MiSeq platforms. Sci Rep 7: 3935.

González-Ramos D, Gorter de Vries AR, Grijseels SS et al. (2016) A new laboratory evolution approach to select for constitutive acetic acid tolerance in *Saccharomyces cerevisiae* and identification of causal mutations. Biotechnol Biofuels 9: 173.

Goodwin S, Gurtowski J, Ethe-Sayers S et al. (2015) Oxford Nanopore sequencing, hybrid error correction, and de novo assembly of a eukaryotic genome. Genome Res 25: 1750–6.

Gorter de Vries AR, Pronk JT, Daran J-MG (2017) Industrial relevance of chromosomal copy number variation in *Saccharomyces* yeasts. Appl Environ Microbiol 83: e03206–16.

Holt C, Yandell M (2011) MAKER2: an annotation pipeline and genome-database management tool for second-generation genome projects. BMC Bioinformatics 12: 491.

Istace B, Friedrich A, d’Agata L et al. (2017) *De novo* assembly and population genomic survey of natural yeast isolates with the Oxford Nanopore MinION sequencer. Gigascience 6: 1–13.

Jansen H, Dirks RP, Liem M et al. (2017) *De novo* whole-genome assembly of a wild type yeast isolate using nanopore sequencing. F1000Research 6.

Kim JM, Vanguri S, Boeke JD et al. (1998) Transposable elements and genome organization: a comprehensive survey of retrotransposons revealed by the complete *Saccharomyces cerevisiae* genome sequence. Genome Res 8: 464–78.

Koren S, Walenz BP, Berlin K et al. (2017) Canu: scalable and accurate long-read assembly via adaptive k-mer weighting and repeat separation. Genome Res 27: 722–36.

Kurtz S, Phillippy A, Delcher AL et al. (2004) Versatile and open software for comparing large genomes. Genome Biol 5: R12.

Li H, Durbin R (2010) Fast and accurate long-read alignment with Burrows-Wheeler transform. Bioinformatics 26: 589–95.

Li H, Handsaker B, Wysoker A et al. (2009) The Sequence Alignment/Map format and SAMtools. Bioinformatics 25: 2078–9.

Lodolo EJ, Kock JLF, Axcell BC et al. (2008) The yeast *Saccharomyces cerevisiae* -the main character in beer brewing. Fems Yeast Res 8: 1018–36.

Loman NJ, Quick J, Simpson JT (2015) A complete bacterial genome assembled *de novo* using only nanopore sequencing data. Nat Methods 12: 733–U51.

Loman NJ, Quinlan AR (2014) Poretools: a toolkit for analyzing nanopore sequence data. Bioinformatics 30: 3399–401.

Loose M, Malla S, Stout M (2016) Real-time selective sequencing using nanopore technology. Nat Methods 13: 751.

Mardis ER (2008) The impact of next-generation sequencing technology on genetics. Trends in genetics: TIG 24: 133–41.

Matheson K, Parsons L, Gammie A (2017) Whole-genome sequence and variant analysis of W303, a widely-used strain of *Saccharomyces cerevisiae*. G3, DOI 10.1534/g3.117.040022g3.117.040022.

McIlwain SJ, Peris D, Sardi M et al. (2016) Genome sequence and analysis of a stress-tolerant, wild-derived strain of *Saccharomyces cerevisiae* used in biofuels research. G3 6: 1757–66.

Milne I, Stephen G, Bayer M et al. (2012) Using Tablet for visual exploration of second-generation sequencing data. Brief Bioinform bbs012.

Naseeb S, Carter Z, Minnis D et al. (2016) Widespread Impact of Chromosomal Inversions on Gene Expression Uncovers Robustness via Phenotypic Buffering. Mol Biol Evol 33: 1679–96.

Naumov GI, Naumova ES, Louis EJ (1995) Genetic mapping of the α-galactosidase *MEL* gene family on right and left telomeres of *Saccharomyces cerevisiae*. Yeast 11: 481–3.

Ng PC, Kirkness EF (2010) Whole genome sequencing. Genetic variation,p.^pp. 215–26. Springer.

Nijkamp J, Winterbach W, Van den Broek M et al. (2010) Integrating genome assemblies with MAIA. Bioinformatics 26: i433–i9.

Nijkamp JF, van den Broek M, Datema E et al. (2012) De novo sequencing, assembly and analysis of the genome of the laboratory strain *Saccharomyces cerevisiae* CEN.PK113-7D, a model for modern industrial biotechnology. Microb Cell Fact 11: 36.

Oud B, Maris AJA, Daran JM et al. (2012) Genome-wide analytical approaches for reverse metabolic engineering of industrially relevant phenotypes in yeast. Fems Yeast Res 12: 183–96.

Papapetridis I, Dijk M, Maris AJA et al. (2017) Metabolic engineering strategies for optimizing acetate reduction, ethanol yield and osmotolerance in *Saccharomyces cerevisiae*. Biotechnol Biofuels 10: 107.

Pryde FE, Huckle TC, Louis EJ (1995) Sequence analysis of the right end of chromosome XV in *Saccharomyces cerevisiae*: an insight into the structural and functional significance of sub-telomeric repeat sequences. Yeast 11: 371–82.

Sović I, Šikić M, Wilm A et al. (2016) Fast and sensitive mapping of nanopore sequencing reads with GraphMap. Nat Commun 7.

Teste M-A, François JM, Parrou J-L (2010) Characterization of a new multigene family encoding isomaltases in the yeast *Saccharomyces cerevisiae*, the *IMA* family. J Biol Chem 285: 26815–24.

Teunissen AW, Steensma HY (1995) Review: The dominant flocculation genes of *Saccharomyces cerevisiae* constitute a new subtelomeric gene family. Yeast 11: 1001–13.

van Dijk EL, Auger H, Jaszczyszyn Y et al. (2014) Ten years of next-generation sequencing technology. Trends in genetics: TIG 30: 418–26.

Venema J, Tollervey D (1999) Ribosome synthesis in *Saccharomyces cerevisiae*. Annu Rev Genet 33: 261–311.

Walker BJ, Abeel T, Shea T et al. (2014) Pilon: an integrated tool for comprehensive microbial variant detection and genome assembly improvement. PLoS One 9: e112963.

Wolfe KH, Shields DC (1997) Molecular evidence for an ancient duplication of the entire yeast genome. Nature 387: 708.

Zhang Z, Schwartz S, Wagner L et al. (2000) A greedy algorithm for aligning DNA sequences. Journal of computational biology: a journal of computational molecular cell biology 7: 203–14.

